# fastCNV: Fast and accurate copy number variation prediction from High-Definition Spatial Transcriptomics and scRNA-Seq Data

**DOI:** 10.1101/2025.10.22.683855

**Authors:** Gadea Cabrejas, Marine Sroussi, Hugo Croizer, Antoine Cazelles, Nicolas Salaün, Lara Jerman, Théo Z Hirsch, Sophie Mouillet-Richard, Pierre Laurent-Puig, Clarice Groeneveld, Aurélien de Reyniès

## Abstract

**Background:** Predicting DNA copy number variations (CNVs) from spatial transcriptomics (ST), including Visium HD, or single-cell RNA-sequencing (scRNA-seq) data helps to distinguish malignant from non-malignant cells and to characterize the clonal architecture of tumor cells. Though there are existing methods of CNV analysis, they are often limited by slow speed, high memory consumption, lower accuracy in the absence of a reference for diploid cells, lower sensitivity at low read counts, and no support for clonal tree construction.

**Results:** To overcome these issues, we developed the R package fastCNV for detecting CNVs from ST, including Visium HD, or scRNA-seq data. FastCNV pools diploid references across samples and, within each sample, aggregates similar spots or cells with few reads into meta spots or cells. It automatically builds a clonality tree, and runs several times faster than other methods while using less memory. To measure the accuracy of fastCNV, we used 117 cancer cell line samples with both scRNA-seq and bulk whole-exome sequencing (WES) data. FastCNV identified CNVs highly correlated to those calculated from WES data (median correlation above 0.75), showing a significant improvement as compared to other methods such as inferCNV. Notably, fastCNV enables, for the first time, the analysis of CNVs from the Visium HD spatial transcriptomics technology. Applied to Visium HD breast cancer ST data, fastCNV identifies tumor subclones tightly related to different histologies, linking specific genetic aberrations to tumor progression.

**Conclusions:** FastCNV is a significant improvement on existing R methods for CNV detection from ST, including Visium HD, or scRNA-seq data in terms of speed, memory usage, sensitivity and accuracy. This highlights its potential to advance cancer research and personalized medicine. FastCNV is available at https://github.com/must-bioinfo/fastCNV/.

## Background

DNA copy number variations (CNVs) play a significant role in diverse pathologies, particularly in cancer. Structural alterations can be pathogenic through deletions inactivating tumor suppressor genes such as *TP53*[1] and *RB1*[2], or through amplifications activating oncogenes such as *MYC*[3] and *ERBB2 (HER2)*[4]. CNVs in cancer progressively accumulate, causing intratumoral heterogeneity and genomic instability. They have been found to be involved in tumor growth[5][6] and drug resistance[7], thus affecting the prognosis of cancer patients[8]. In a clinical context, accurate detection of CNVs is essential to inform personalized therapeutic strategies[9][10]. This detection is typically performed through DNA sequencing approaches, such as whole-exome sequencing (WES) or whole genome sequencing (WGS), assisted by computational tools including ControlFREEC[11], Facet[12] or Wisecondor[13].

Several computational tools have been developed to infer CNVs from RNAseq data. They are based on the assumption that CNVs directly affect transcript abundance, in a way that amplifications usually lead to overexpression and deletions lead to underexpression[14][15]. The relative level of expression against a diploid reference is used to predict losses and gains from patterns of over- and underexpression[15]. Nevertheless, not every one of the CNVs predicted from transcriptomic data translates into actual genomic alterations since gene expression is also determined by epigenetic mechanisms. Consequently, RNA-based CNV inference lacks the specificity required for clinical diagnostics. Nonetheless, it is extremely helpful in a research context. Notably, the inference of putative CNVs from single-cell or spatial transcriptomic datasets offers valuable insights into tumor heterogeneity and clonal architecture at less cost and complexity than parallel DNA-based profiling. Given the current technical and financial constraints on high-resolution DNA sequencing at the single cell or spatial level, transcriptome-based inference of CNVs represents a scalable and cost-effective alternative for studying tumor evolution and cellular diversity in situ[16].

Several methods have been developed to infer CNVs from ST or scRNA-seq data, including inferCNV[17], Numbat[18], CopyKAT[19], CaSpER[20], SCEVAN[21], SpatialInferCNV[22], CONICSmat[23], stmut[24], HoneyBADGER[25]. Among these methods, inferCNV is currently the most widely adopted tool within the R community (Supplementary Table 1). Numbat stands out for its haplotype-aware inference, which enables the detection of copy-neutral loss of heterozygosity (cnLOH). However, Numbat reliance on raw sequencing data (FASTQ/BAM) limits its applicability on public datasets when only processed gene expression matrices are available.

**Supplementary Table 1 :**
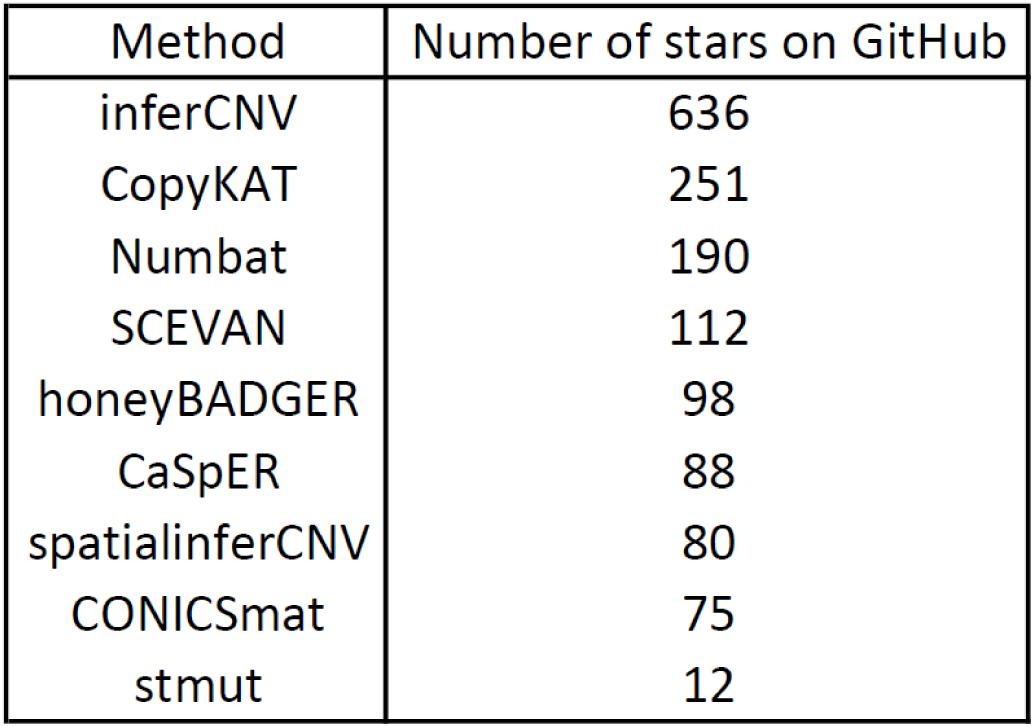
Github popularity of tools for inferring CNVs in transcriptomics data Number of GitHub stars (as of September 2025) for selected tools used to infer copy number variation (CNV) from single-cell or spatial transcriptomic data.

Our primary motivation for developing fastCNV stemmed from the challenges we encountered with existing R-based tools, which are too slow and insufficiently flexible to seamlessly handle either ST or scRNAseq data indifferently[26][27]. fastCNV was designed to address these challenges, offering improved computational efficiency and compatibility with diverse transcriptomic modalities. In addition, it incorporated advanced features such as integrated visualization tools and automated clonality tree construction, enabling comprehensive exploration of tumor heterogeneity and evolution.

To assess the accuracy of fastCNV results, we applied it to the scRNA-seq data of 117 cancer cell lines published by Kinker *et al*[28]. Bulk WES data were also available for these 117 cell lines, all included in the Cancer Cell Line Encyclopedia (CCLE) database. The WES-based CNV profiles of these 117 cell lines were considered here as a ground-truth reference to benchmark fastCNV results. We further demonstrated the applicability of fastCNV to ST data by showing that it accurately discriminates tumor from non-tumor regions, using histopathological annotations as a reference standard. Importantly, we observed that distinct CNV-derived clones often correspond to regions with different morphological features within the tumor, highlighting the biological relevance of the inferred clonal architecture. Finally, we tested fastCNV on a Visium HD dataset, demonstrating its ability to successfully process high-resolution spatial transcriptomic data. To our knowledge, this constitutes the first CNV analysis performed on Visium HD, further underscoring the versatility of the fastCNV method.

## Implementation

### Input data

The fastCNV R package processes spatial transcriptomics or single cell RNA-sequencing data, previously imported using the Seurat R package [29] which produces Seurat objects. The main input of the *fastCNV* function is either a Seurat object, when dealing with one sample, or a list of Seurat objects, when dealing with several samples **(Fig. 1)**. Working with a list of Seurat objects (each corresponding to one sample) allows us to share the information of several samples for some analytical steps, as illustrated later.

**Figure 1:**
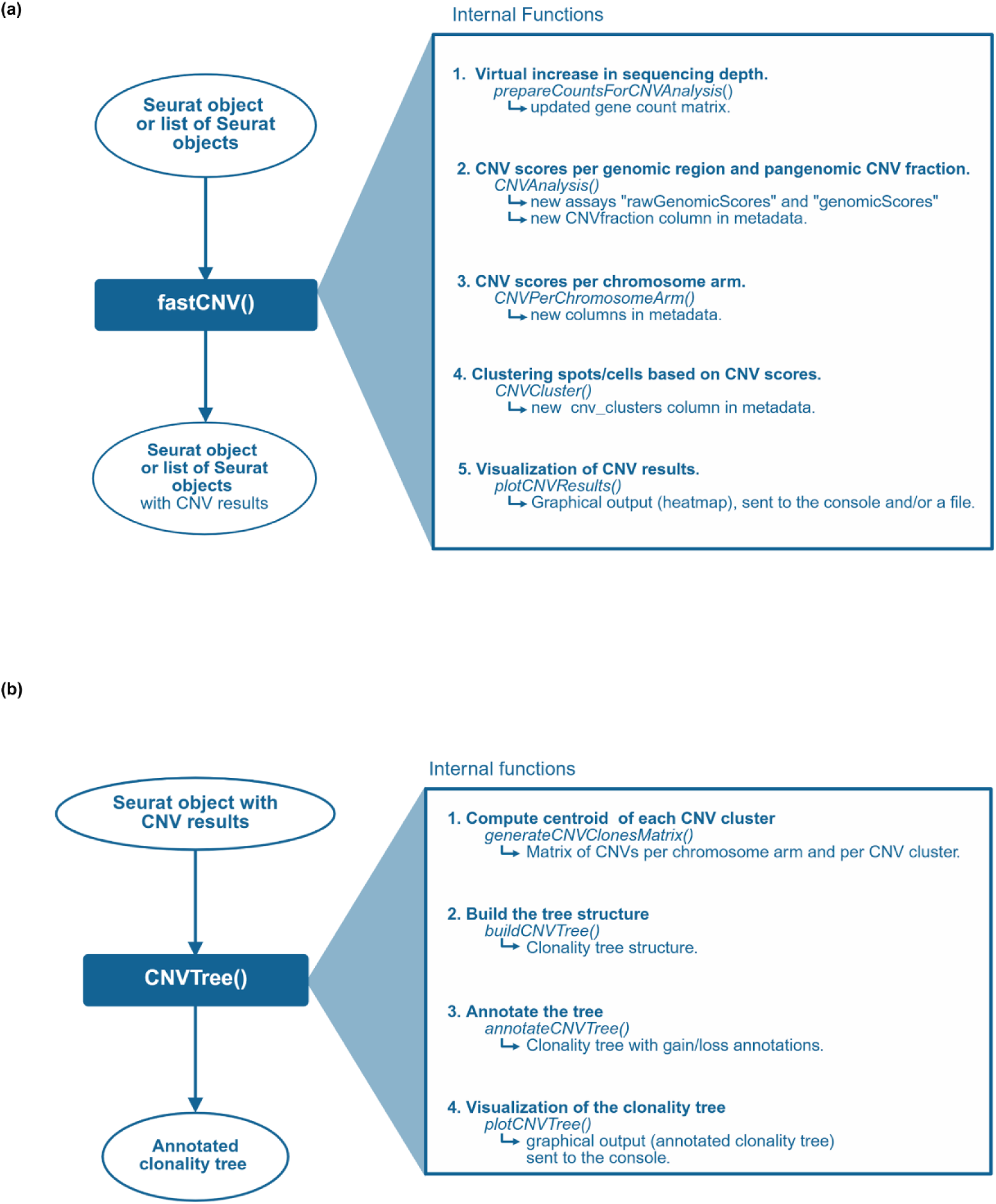
Overview of the fastCNV workflow **(a**) The fastCNV() function takes as input a Seurat object (or a list of Seurat objects) and processes it to predict CNVs. It returns a Seurat object (or list of Seurat objects) with the corresponding CNV results. Optionally a graphical output (pangenomic heatmap(s)) can also be produced. **(b)** The CNVTree() function takes as input a Seurat Object with CNV results and returns an annotated clonality tree and its graphical representation.

### Diploid reference

Specifying spots/cells corresponding to a diploid reference is expected to improve the accuracy of fastCNV results. This can be done by indicating a variable (from the metadata table) and the related values corresponding to a diploid status. For example, for a variable called ‘cell type’ with the following values: ‘tumour’, ‘normal epithelium’, ‘immune’ and ‘stromal’, values other than ‘tumour’ would typically correspond to a diploid status. If this information is provided, fastCNV will gather the diploid reference spots/cells from all Seurat objects, meaning that the same reference will be used to analyze all Seurat objects. This pooling approach supposes that samples are of similar types, as it appears very challenging to define a universal diploid reference (for transcriptome data).

If no diploid reference is provided, which might produce suboptimal CNV results, all spots/cells will serve as the “diploid” reference and their pooling across Seurat objects is then optional.

### Virtual increase in sequencing depth

A low sequencing depth, resulting in a low number of reads per spot/cell, is a limiting factor for the identification of CNVs. As a consequence, fastCNV first proceeds with a step aimed at aggregating similar spots/cells in meta-spots/meta-cells, in order to reach a sufficient total number of reads for the identification of CNVs (Cf. *prepareCountsForCNVAnalysis* function). To identify similar spots/cells, we perform unsupervised Louvain clustering of spots/cells, based on the gene count matrix. The obtained clusters of spots/cells are further subdivided by cell type or histological feature, if this information is given. Within each (sub)cluster, the idea is now to obtain any partition (i.e., binning) of the spots/cells allowing to aggregate them into meta-spots/cells, each containing a minimum total number of reads (by default: 15,000 reads). We simply build this partition sequentially. Typically, in Visium (non-HD) ST data, meta-spots correspond to 1 to 3 original spots, using default parameters. This step leads to a new gene count matrix at the level of meta-spots or meta-cells. Finally, in the original gene count matrix, we impute each spot/cell with the corresponding meta-spot/meta-cell. Doing so we keep the original granularity in terms of spots/cells, while getting more reads per spot/cell.

### Genomic regions

To calculate CNV scores per genomic region (Cf. *CNVAnalysis* function), we need to define genomic regions, i.e. segments of the genome including a certain number of genes.

We first map all the genes present in the gene count matrix (of the Seurat objects) to the human genome, using the gene information downloaded from Ensembl[30], which provides for each gene the corresponding chromosome and start/end positions. We also determine and store for each gene the chromosome arm information. Genes are then ordered according to their chromosome numbers and start positions.

As genes with low or null expression across most spots/cells are expected to be less informative for CNV identification, we keep only the N most expressed genes (default N=7000), based on the average of raw counts across spots/cells from the diploid reference. If in any given chromosome arm, less than 200 genes are among the N selected genes, the 200 most expressed genes in that chromosome arm are also kept ; in this case, the final gene selection contains more than N genes.

The above selected genes (ordered by genomic positions) are then used to build overlapping genomic regions covering all the genome, by using a moving window per chromosome arm and based on two user-defined parameters : (i) the size of each region, i.e. the number of genes it shall contain (default: 150 genes), and (ii) the step between two consecutive regions (default : 10 genes). Using the default parameters, about 250 genomic regions are obtained.

### Raw CNV scores

The gene count matrix is log2-transformed (after adding 1 to all values) then zero-centered per spot/cell, yielding a normalized gene expression matrix. We then calculate per gene a scaling factor, corresponding to the average normalized expression across spots/cells serving as the diploid reference. When the reference is defined by several modalities of a variable (ex. cell type in {“normal epithelium”, “immune”, “stromal”}) the scaling factor is calculated as the average of the averages per modality. For each gene, the normalized data are scaled by subtracting the corresponding scaling factor. Then values below −3 are set to −3, and those above 3 are set to 3; this truncation is performed to avoid overweighting genes with a high dynamic range. This yields a scaled normalized gene expression matrix, with all values between −3 and 3.

For each genomic region, raw CNV scores are calculated per spot/cell as the average of the above defined scaled normalized data, across all genes included in the genomic region. We thus obtain a (genomic regions x spots/cells) matrix of CNV scores, for each Seurat object. This matrix is added to each Seurat object, as a new assay, labelled “rawGenomicScores”. By construction, all CNV scores range from −3 to 3. CNV scores around zero generally correspond to a diploid status, while negative and positive CNV scores typically indicate losses and gains, respectively.

### Denoised CNV scores

From the raw CNV scores we derive denoised CNV scores, where raw CNV scores around zero are set to zero. More precisely, we calculate per genomic region the 1st and 99th percentiles of raw CNV scores across spots/cells serving as the diploid reference. CNV scores falling between these two percentiles are then set to zero. The resulting matrix is added to each Seurat object, as a new assay, labelled “genomicScores”. The denoised CNV scores enhance the contrast between (putative) diploid and non-diploid statuses, which can notably be useful for graphical visualizations.

### CNV fractions

The pangenomic CNV fraction for each cell/spot is computed as the proportion of non-zero denoised CNV scores across all genomic regions, independently for each Seurat object. The corresponding vector is added to the metadata matrix of each Seurat object, as a new column labelled “cnv_fraction”. The CNV fraction of any spot/cell ranges from 0 to 1, and reflects the proportion of genomic regions containing CNVs for that spot/cell, that is a score of chromosomal instability (CIN score). For a spot (ST data), it might also reflect the tumor purity (i.e., the proportion of tumor non-diploid cells within that spot).

### CNV scores per chromosome arm

FastCNV computes CNV scores per chromosome arm (using denoised scores by default), resulting in a (spots/cells x chromosome arms) matrix (Cf. *CNVPerChromosomeArm* function). Each cell of this matrix contains the average CNV score across the genomic regions of the related chromosome arm, for the concerned spot/cell. This information is inserted as new columns (one per chromosome arm) in the metadata table of each Seurat object.

### Clustering of spots/cells based on CNV scores

Within each Seurat object, we cluster spots or cells based on their CNV scores (using denoised scores by default), thus enabling subclonality to be detected, with distinct clusters assumed to represent distinct subclones (Cf. *CNVCluster* function). This is performed using hierarchical clustering, with Ward linkage and Manhattan distance. The optimal number of clusters is automatically determined by identifying the inflection point, known as the elbow point, on the within-cluster sum of squares curve. Sometimes we obtain highly similar clusters differing solely slightly in terms of CNV scores intensities, which means that the same subclone is separated into several clusters. To avoid this problem, fastCNV merges clusters whose centroids are highly correlated.

### Pangenomic visualization of CNVs

The *plotCNVResults* function produces a pangenomic heatmap visualization of the CNV scores, for a given Seurat object. It is based on the complexHeatmap[31] package. Users can easily modify the annotations plotted in the heatmap, print it to the R console, and/or save it as a PDF or PNG file.

### Clonality tree

The *CNVTree* function allows building a clonality tree from a Seurat object that includes CNV results. It starts with the (CNV-based) clusters of spots/cells, seen as subclones, and computes the centroid of each cluster. It thus produces a matrix (genomic regions x clusters) containing the average CNV score across spots/cells of each cluster for each genomic region (cf. *generateCNVClonesMatrix* function). It then computes pairwise distances between the cluster’s centroids, using the euclidean distance by default. The resulting distance matrix is then analysed with phylogenetic functions from the ape[32] or phangorn[33] R packages (depending on the options chosen by the user) to infer the clonality tree structure (Cf. *buildCNVTree* function).

Each branch of the clonality tree is then automatically annotated in terms of the chromosome arm gains and losses shared by all children branches and not present in the parent branch (Cf. *annotateCNVTree* function). A user-defined threshold (to be applied on CNV scores) is used to filter the CNV events (i.e., gains and losses) considered in this analysis. The annotations (CNV events) of the clonality tree can also be manually curated to refine the final output.

### Visualization of the clonality tree

The annotated clonality tree can be plotted using the *plotCNVTree* function, which is based on the ggtree package[34], enabling various customizations.

## Results

### Validation of fastCNV on CCLE cell line data

To evaluate the reliability of fastCNV results, we used 117 cancer cell lines from the CCLE database [35], for which both bulk WES and scRNA-seq data were publicly available. We used the WES-based CNV data (gene level log copy number) as the ground-truth reference. It was retrieved using the ExperimentHub R package (experiment ID: EH7555). The scRNA-seq data (raw gene counts) of these cell lines included 41035 cells and were downloaded from the Broad Institute’s single cell portal (series ID: SCP542).

We applied fastCNV to the scRNA-seq data of each cell line, obtaining raw and denoised CNV scores per genomic region and per cell. We did not specify a diploid reference in that case, as it was difficult to define given the wide diversity of cancer types in this series. We averaged scores across all cells of each cell line, obtaining a (genomic regions x cell lines) matrix for the raw CNV scores, and another for the denoised ones. We also averaged, per genomic region, the gene level WES-derived CNV data, obtaining a (genomic regions x cell lines) matrix. We then calculated per cell line the correlation between CNV scores obtained from fastCNV and those coming from WES data, as illustrated for the cell line TE10_OESOPHAGUS (**Fig. 2a**). FastCNV scores were overall highly correlated to the ground-truth, raw CNV scores showing a higher correlation to the reference (median=0.76, mean 0.72) than denoised CNV scores (median=0.71, mean=0.67) (**Fig. 2b**). The few cell lines with a weak correlation between WES-based (ground truth) and scRNA-seq derived CNV scores mostly had little to no CNV (**Fig. 2c**), those having a correlation below 0.5 showing a much lower CNV fraction (CIN score) than the other cell lines (Welch T-test p < 0.001). The cell lines with a low (<10%) WES-based CNV fraction largely corresponded to those having a low fastCNV-based CNV fraction (<10%) (Chi2 test pvalue < 1e-10), showing that fastCNV identifies accurately tumor with a low CNV fraction.

**Figure 2:**
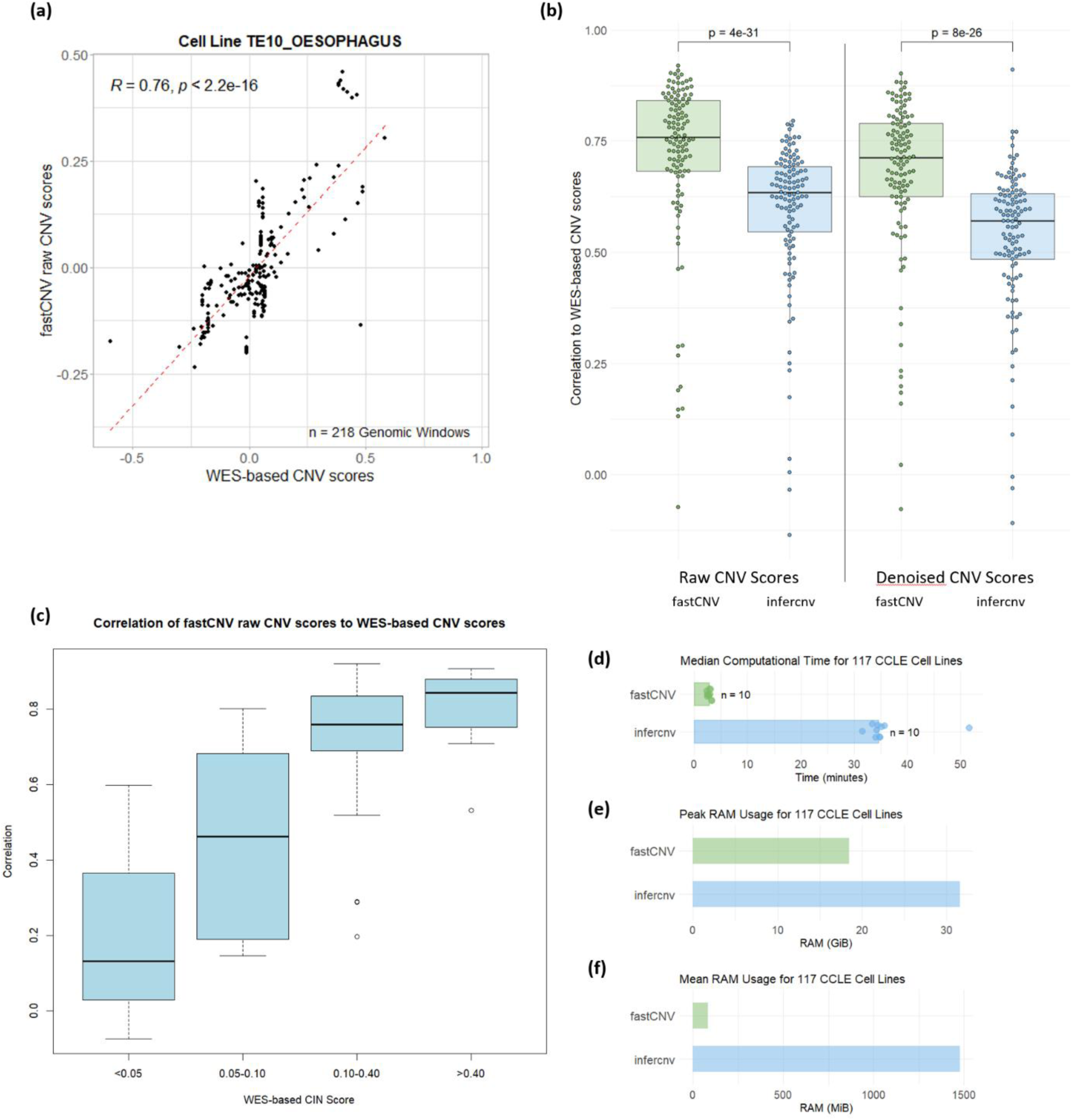
Validation of CNV scores on scRNA-seq data of CCLE Cell Lines **(a)** Scatter plot comparing raw CNV scores obtained using fastCNV on scRNA-seq data with WES-based CNV scores, for the TE10_OESOPHAGUS cell line. The dashed line represents the corresponding linear model. **(b)** Boxplot showing the distribution of correlations between predicted CNV scores using fastCNV (green) and inferCNV (blue) on scRNA-seq data (raw scores on the left, denoised scores on the right) and WES-based CNV scores, for 117 CCLE cell lines (each dot corresponds to a cell line). Pvalues correspond to paired T-tests comparing fastCNV and inferCNV respective correlations to WES-based CNVs. **(c)** Boxplot showing the correlation between fastCNV raw CNV scores (scRNA-seq data) and WES-based CNV scores across 117 CCLE cell lines, grouped by degree of WES-based CNV fraction (<5%, 5-10%, 10-40%, >40%). **(d)** Barplot showing the median computational time for fastCNV and inferCNV on 117 cell lines. **(e)** Barplot showing the peak RAM usage for fastCNV and inferCNV on 117 cell lines. **(f)** Barplot showing the mean RAM usage for fastCNV and inferCNV on 117 cell lines.

Overall, these results demonstrate, across a broad range of cancer cell lines, that fastCNV provides accurate CNV results.

### Comparison of fastCNV and inferCNV on CCLE cell line data

We calculated raw and denoised CNV scores using the inferCNV package on the same scRNA-seq data. For each cell line, the output of inferCNV included a (genes x cells) matrix of raw CNV scores, and another with denoised CNV scores. This output is not directly referring to genomic regions, but rather to a set of about 2000 genes along the genome. However, the score for any of these genes is obtained by aggregating information from many genes around, so here a gene represents *de facto* a genomic region. We averaged these scores per cell line, obtaining two (genes x cell lines) matrices, with averaged raw and denoised scores respectively. We then extracted from the WES-based CNV data the subpart corresponding to the genes in common with the inferCNV output, getting a (genes x cell lines) matrix. We then calculated the correlation per cell line between inferCNV scores and WES-based scores. Here again raw scores were more correlated to the ground-truth (median=0.63, mean=0.59) than denoised ones (median=0.57, mean=0.53) (**Fig. 2b**).

The CNV scores calculated using fastCNV were significantly more correlated to the ground truth than those calculated using inferCNV, both for raw and denoised scores (paired T-test p values less than 4e-31 and 8e-24 respectively, **Fig. 2b**), clearly showing that fastCNV outperforms inferCNV in terms of accuracy. Regarding computational efficiency (**Fig. 2d**), fastCNV was approximately 10 times faster than inferCNV, completing CNV inference for the 117 cell lines in around 4 minutes, compared to 40 minutes for inferCNV. Here, the mean RAM usage of fastCNV was around 6% that of inferCNV (80 MiB versus 1400 MiB, **Fig. 2e**) and the peak RAM usage of fastCNV was 60% that of inferCNV (19 GiB versus 32 GiB, **Fig. 2f**).

### Application of fastCNV to colon cancer scRNA-seq data

We applied fastCNV to a colon cancer scRNA-seq sample (ID C113) from Pelka *et al.* series [36] (**Fig. 3**), containing epithelial, lymphoid, myeloid and mesenchymal cells (for a total of 1454 cells). We used non-epithelial cells as the diploid reference. The matrix of denoised CNV scores is shown as a pangenomic heatmap, with genomic regions (columns) ordered by chromosome, and cells (rows) organized in four CNV-based clusters (**Fig. 3a**). FastCNV infers a clonality tree from centroids (at the chromosome arm level) of these four clusters (**Fig. 3b**). The CNV annotations of the tree branches are derived from these centroids, discretized using a default threshold (**Fig. 3c**). To improve the CNV annotations of the clonality tree we manually curated them (**Fig. 3 d-e**). We then calculated the UMAP embedding of the gene count matrix, thus identifying nine groups of cells, clearly related to distinct cell lineages, with epithelial (n=4), lymphoid (n=3), myeloid (n=1), and mesenchymal (n=1) groups (**Fig. 3f**). The four epithelial groups map to distinct subclones (**Fig. 3g**) and show different CNV fractions (**Fig. 3h**), three of them being clearly malignant. One of the epithelial groups shows a very low CNV fraction, and might correspond to pre-malignant cells. In conclusion, fastCNV allows here to identify malignant cells and the related clonal architecture, explaining a large part of the groups of cells that emerge from unsupervised analysis through UMAP embedding.

**Figure 3:**
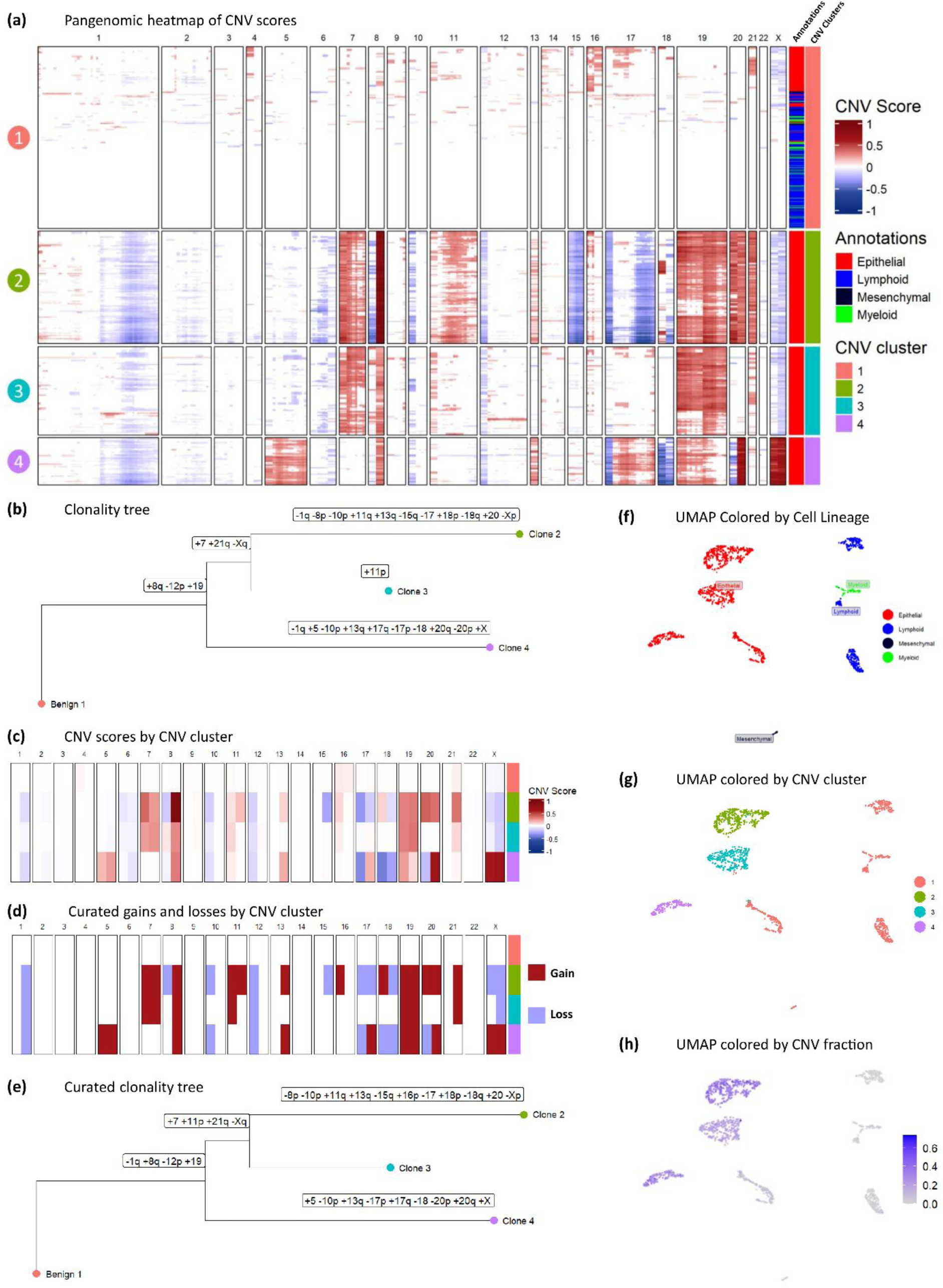
Illustration of subclonal heterogeneity in scRNA-seq colon cancer data Analysis of the C113 colon cancer scRNA-seq sample from Pelka *et al* series. **(a)** Pangenomic heatmap showing the (cells x genomic regions) matrix of fastCNV denoised CNV scores. The genomic regions are in genomic order, the cells are ordered according to CNV clusters (1 to 4). CNV scores corresponding to copy gains (respectively losses) are in red (respectively blue), those corresponding to a diploid status are in white. The cell lineage of each cell is indicated (red: epithelial, blue: lymphoid, black: mesenchymal, green: myeloid). **(b)** Phylogenetic tree based on CNV clusters 1 to 4, seen as subclones, depicting the corresponding subclonal evolution in terms of CNV events. **(c)** Heatmap showing CNV scores per CNV cluster and chromosome arm, computed from the centroid profiles of each CNV cluster. **(d)** Heatmap displaying discretized CNV states by chromosome arm (copy gain: red, copy loss: blue, diploid: white) across CNV clusters after curation. **(e)** Phylogenetic tree with curated CNV annotations along the branches. **(f)** UMAP plot of the normalized scRNA-seq data showing cells colored by their lineage. **(g)** UMAP plot of single cells colored by CNV-based clusters. **(h)** UMAP plot showing the fraction of CNV per cell.

### Application of fastCNV to colon cancer Visium ST data

We applied fastCNV to a colon cancer Visium ST FFPE sample (10x Genomics) from Valdeolivas *el al* series [36], containing spots annotated as tumor, non-tumor mucosa and immune cells (**Fig. 4a-b**). All the spots, except the tumor ones, were used as the diploid reference. The denoised CNV scores are presented in the form of a pangenomic heatmap, with spots organized in three CNV-based clusters (**Fig. 4c**). The centroids of these three clusters were used to infer a clonality tree and to annotate it (**Fig. S1**). To improve the CNV annotations of this clonality tree we manually curated them (**Fig. 4 d-e**). The subclones are spatially segregated: subclone 1, devoid of CNV, maps to the non-tumor region, subclone 2, with numerous CNVs, maps to the right part of the tumor region and subclone 3, with an intermediate number of CNVs, maps to the left and bottom part of the tumor region (**Fig. 4f**). Accordingly, the spatial distribution of the CNV fractions also shows clear regional patterns (**Fig. 4g**).

**Figure 4:**
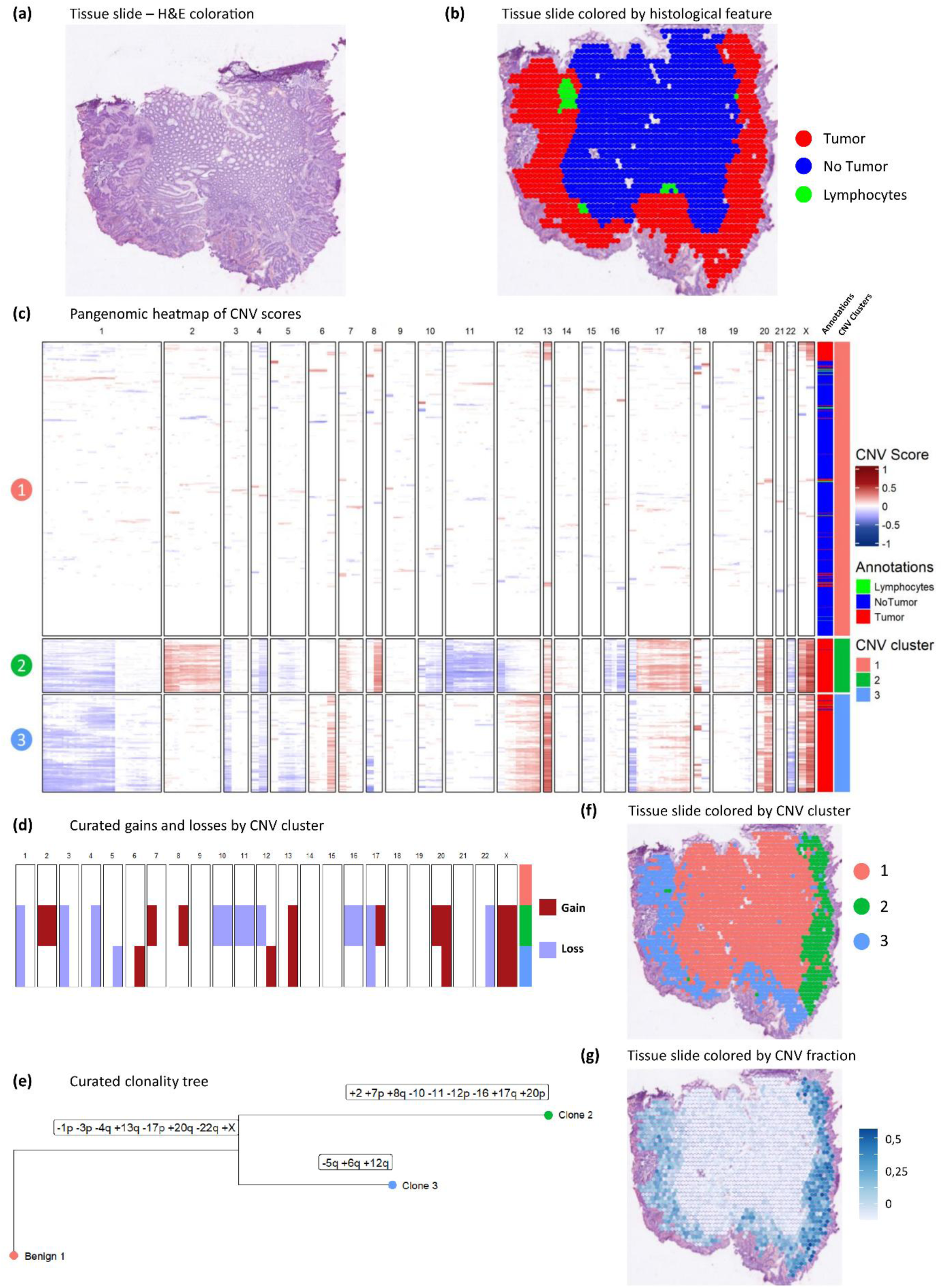
Illustration of subclonal heterogeneity in Spatial Transcriptomics colon cancer data. Analysis of the SN048_A121573_Rep1 colon cancer Visium ST sample from Valdeolivas *et al* series. **(a)** H&E coloration of the tissue slide. **(b)** Spots of the Visium ST slide are colored according to the histological annotation (tumor: red, non-tumor: blue, immune cells: green). **(c)** Pangenomic heatmap showing the (spots x genomic regions) matrix of fastCNV denoised CNV scores, with spots ordered according to CNV clusters (1 to 3). The histological annotations per spot are given on the right. **(d)** Heatmap displaying discretized CNV events by chromosome arm (copy gain: red, copy loss: blue, diploid status: white) across CNV clusters after curation. **(e)** Phylogenetic tree based on CNV clusters 1 to 3. Branches are annotated using curated CNV events. Spots of the Visium ST slide are colored according to **(f)** CNV clusters (1: red, 2: green, 3: blue) and **(g)** pangenomic CNV fraction (low: white, high: blue).

**Supplementary Figure 1 :**
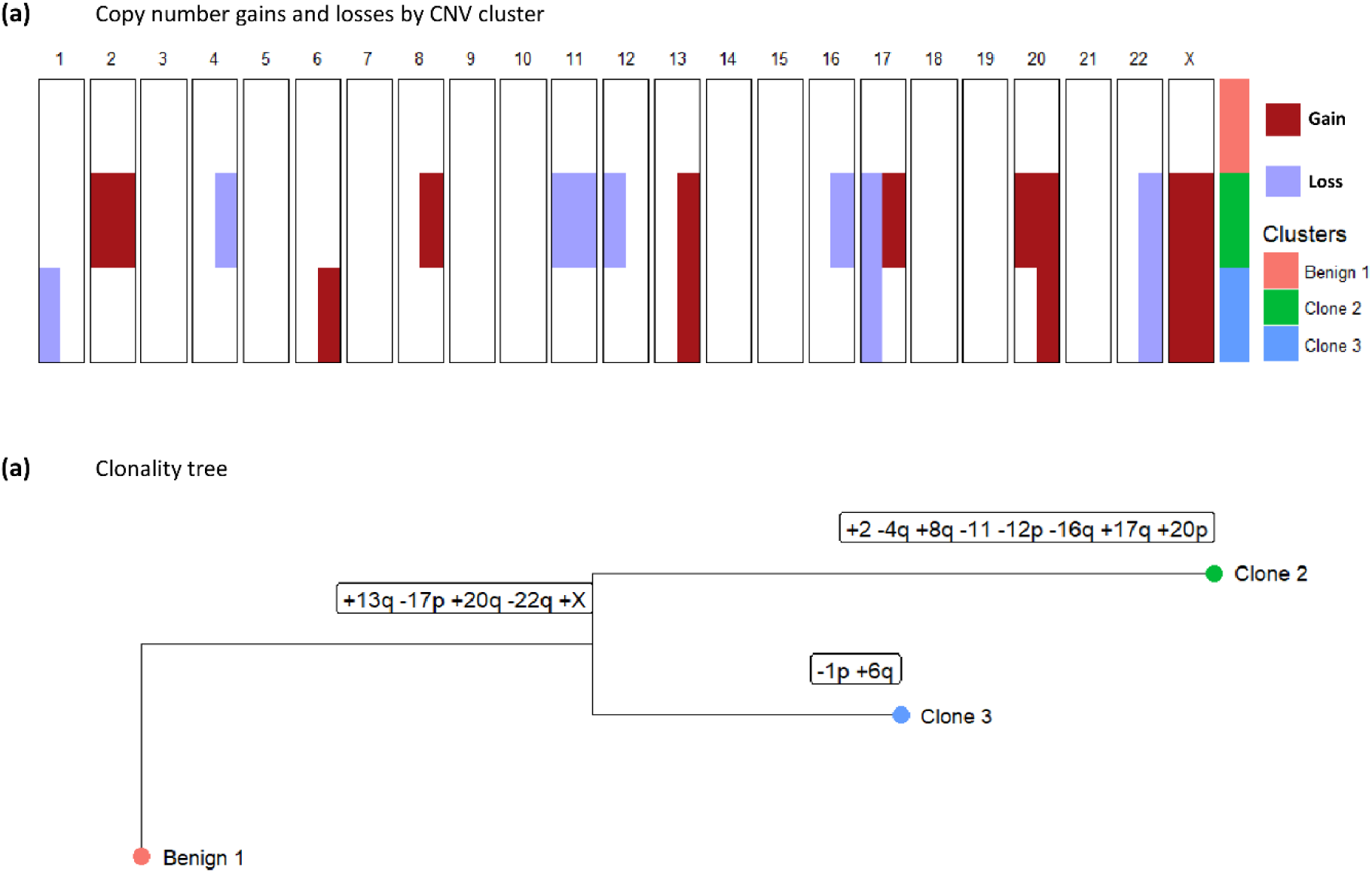
Illustration of subclonal heterogenetity prior to CNV curation on colon cancer ST sample. Analysis of the SN048_A121573_Rep1 colon cancer Visium ST sample from Valdeolivas *et al* series. **(a)** Heatmap displaying discretized CNV events per chromosome arm (copy gain: red, copy loss: blue, diploid status: white) across CNV clusters (1 to 3) before curation. **(b)** Phylogenetic tree based on CNV clusters 1 to 3, with branches annotated using discretized chromosome arm level CNV events, before curation.

### Application of fastCNV to breast cancer Visium HD ST data

We applied fastCNV to a breast cancer Visium HD sample obtained from the 10x Genomics website (**Fig. 5**). Despite the high computational demands of this dataset — approximately 500,000 spots at 8 μm resolution and 150,000 spots at 16 μm resolution, fastCNV runs efficiently on a high-memory server, with peak RAM usage of ∼200 GB for 8 μm resolution. At 16 μm resolution, the analysis can also be performed on a local workstation with 64 GB of RAM (peak usage ∼50 GB). This sample contains tumor and non-tumor regions; the latter serves as the diploid reference (**Fig. 5a**). Based on the denoised CNV scores, six CNV-based clusters are identified (**Fig. 5b**). Some of these clusters show very different CNV fractions (**Fig. 5c**) and map to distinct tumor regions (**Fig. 5d**). The spatial distribution of these clusters shows that they corresponded closely to morphological patterns: cluster 4, with extensive CNVs, corresponds to a region of high histological grade, cluster 6 highlights a preinvasive ductal carcinoma in situ (DCIS) lesion, suggesting it evolved independently from the invasive tumor and did not acquire invasive traits, while cluster 5 characterized by the typical 1q gain and minimal rearrangement corresponds to a non-tumoral dysplastic lesion (**Fig. 5e**). Additional CNV metrics, such as CNV fraction and arm-level alterations, also delineate tumor subregions with distinct genomic profiles such as 11q gain or loss (**Fig. 5f-g**). Based on the CNV clusters, fastCNV reconstructed a clonality tree that describes the presumed sequence of subclone emergence and tumor evolution (**Fig. 5h**). Collectively, these results demonstrate that fastCNV captures high-resolution, spatially resolved CNV heterogeneity, offering insights into tumor composition and evolutionary dynamics. To our knowledge, it is the only method demonstrated to run on Visium HD data.

**Figure 5:**
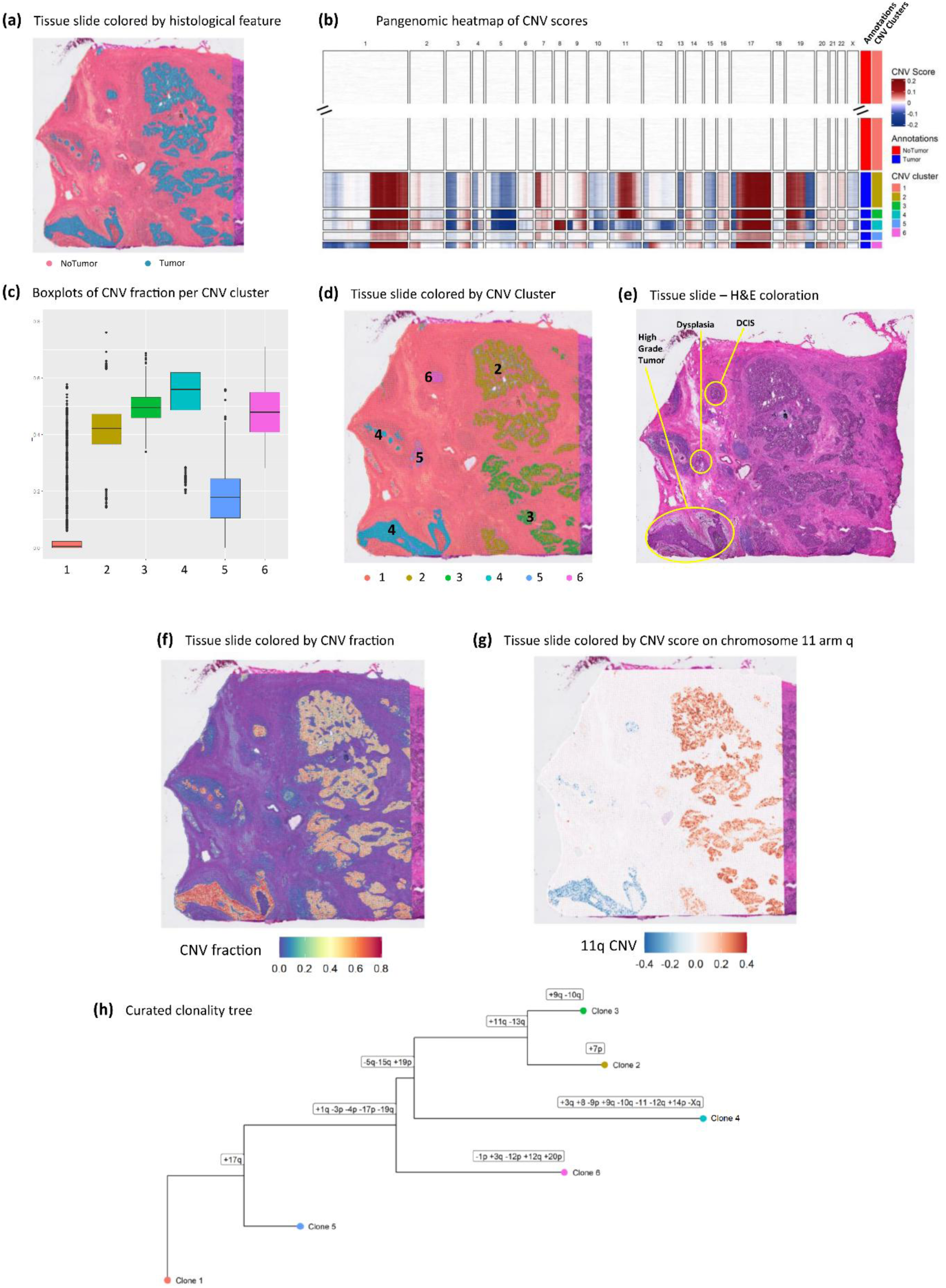
Illustration of subclonal heterogeneity in Visium HD Breast cancer data Analysis of the Visium_HD_FF_Human_Breast_Cancer Visium HD sample from the 10x Genomics database. **(a)** Spots of the Visium HD slide are colored according to tumor status (tumor: blue, non-tumor: pink). **(b)** Pangenomic heatmap showing the (spots x genomic regions) matrix of fastCNV denoised CNV scores, with spots ordered according to CNV clusters (1 to 6). The histological annotations per spot are given on the right. **(c)** Boxplots showing the pangenomic CNV fraction per CNV cluster. **(d)** Spots of the Visium HD slide are colored according to the CNV clusters (1 to 6). **(e)** H&E-stained slide, with contours focusing on regions related to distinct histologies. Spots of the Visium HD slide are colored according to **(f)** the pangenomic CNV fraction and **(g)** the CNV status for chromosome 11 arm q. **(h)** Phylogenetic tree based on CNV clusters 1 to 6. Branches are annotated using curated CNV events at the chromosome arm level.

## Conclusion

fastCNV is a comprehensive tool balancing speed and accuracy to support the analysis of increasingly large and complex transcriptomics datasets, such as those generated by Visium HD technology. With efficient support for high-resolution data at tens to hundreds of thousands of spatial spots, fastCNV addresses the significant need for scalable CNV detection in spatial contexts. Its ability to distinguish tumor from non-tumor regions and resolve subclonal architecture provides valuable insights into tumor heterogeneity. Moreover, fastCNV’s reduced computational resource requirements compared to existing approaches makes it accessible to a wider range of research settings. FastCNV also offers practical advantages in terms of ease of deployment. Unlike inferCNV, which relies on external dependencies such as JAGS that can present installation challenges in certain computing environments, fastCNV has minimal dependency requirements, facilitating straightforward installation. As transcriptomics moves forward towards higher resolution and complexity, fastCNV offers a robust foundation for further research aimed at unraveling the genetic and spatial dynamics of complex tissues.

## Availability and requirements

**Project name :** fastCNV

**Project home page** : https://github.com/must-bioinfo/fastCNV/

**Operating system** : Platform independent

**Programming language** : R

**Other requirements** : Seurat v5

**License** : GPL-3

**Any restrictions to use by non-academic :** None

## List of abbreviations

CCLE: Cancer Cell Line Encyclopedia
cnLOH: copy-neutral loss of heterozygosity
CNV: Copy Number Variations
scRNA-seq: Single Cell RNA Sequencing
ST: Spatial transcriptomics
WES: Whole Exome Sequencing
WGS: Whole Genome Sequencing

## Declarations

### Ethics approval and consent to participate

Not applicable.

### Consent for publication

Not applicable.

### Availability of data and materials

The scRNA-seq data for the 117 cancer cell lines from the CCLE are publicly available via the Broad Institute’s Single Cell Portal under accession number SCP542.

The scRNA-seq colon expression count matrix is accessible through GEO under accession GSE178341.

The spatial transcriptomics slide of colon cancer tissue used in this study is available on Zenodo at https://zenodo.org/records/7760264.

The Visium HD breast slide can be accessed via 10x Genomics’ public datasets portal at https://www.10xgenomics.com/datasets.

The fastCNV R package is available at https://github.com/must-bioinfo/fastCNV/.

## Methods

### Selecting the CCLE cell lines

From the original pool of cell lines in *Kinker et al* (n = 198), we selected those with more than 200 cells (n = 122) passing quality control. Then, we subsetted this list to only those for which the true copy number was available in the R package ExperimentHub[38], resulting in a final set of 117 cell lines.

### Histological annotations for the 10X Visium FFPE colorectal cancer slide[37]

Histological annotations for the 10X Visium FFPE colorectal cancer slide[37] were derived from those provided in the original publication. For improved visibility in the analysis, the original categories were grouped as follows: “epithelium&submucosa”, “non neo epithelium”, and “submucosa” were grouped under “NoTumor”; “IC aggregate_submucosa” was relabeled as “Lymphocytes”; and “tumor”, “tumor&stroma_IC met to high”, and “stroma_fibroblastic_IC high” were combined into the “Tumor” category.

### Versions of the R packages used

R version 4.4.2

Seurat version 5.1.0

SeuratObject version 5.0.2

inferCNV version 1.20.0

fastCNV 1.1.0

fastCNVdata version 1.0.3

ComplexHeatmap version 2.20.0

ape version 5.8

phangorn version 2.12.1

ggtree version 3.12.0

ExperimentHub version 2.12.0

All tests were done in a 20 CPU 64GB RAM machine.

## Author Contributions

**GC** : Development, Data analysis, Interpretation, Writing – original draft, Writing – review & editing.

**MS** : Development, Data Analysis.

**HC** : Data Analysis, Interpretation, Writing – review.

**AC** : Interpretation.

**NS** : Feedback.

**LJ** : Data Analysis.

**TZH** : Data Analysis, Interpretation, Feedback, Writing – review & editing.

**SMR** : Data Analysis, Interpretation, Feedback.

**PLP** : Interpretation, Writing – review & editing.

**CS** : Supervision, Development, Interpretation, Feedback, Writing – review & editing.

**AdR** : Supervision, Development, Interpretation, Feedback, Writing – review & editing.

All authors read and approved the final manuscript.

## Acknowledments

We thank Roseline Vibert, Camilla Pilati, Bastien Rance and Alix Yan for fruitful discussions.

